# Curvature-induced cell rearrangements in biological tissues

**DOI:** 10.1101/2022.05.18.492428

**Authors:** Yuting Lou, Jean-Francois Rupprecht, Tetsuya Hiraiwa, Timothy E Saunders

## Abstract

On a curved surface, epithelial cells can adapt to geometric constraints by tilting and by exchanging their neighbors from apical to basal sides, known as an apicobasal T1 (AB-T1) transition. The relationship between cell tilt, AB-T1 transitions, and tissue curvature still lacks a unified understanding. Here, we propose a general framework for cell packing in curved environments and explain the formation of AB-T1 transitions under different conditions. We find that steep curvature gradients can lead to cell tilting and induce AB-T1 transitions. Conversely, large curvature anisotropy can drive AB-T1 transitions by hydrostatic pressure. The two mechanisms compete to determine the impact of tissue geometry and mechanics on optimized cell rearrangements in 3D.

---

As the external surfaces and barriers of many organs, epithelial tissues have to mechanically adapt to their environment [1, 2]. Extensive research into cell shape in 2D [3–10] and 3D [11–14] has revealed insights into how cells pack and undergo rearrangement during epithelial tissue formation [7–10, 15]. Cellular dynamic processes, like division and apoptosis, can rearrange cell neighbors. T1-transitions - the exchange of neighbors without altering the cell number - is another ubiquitous mechanism of cell rearrangements [16, 17]. T1 transitions are important in mediating planar tissue dynamics. For example, oriented T1 transitions can lead to tissue elongation or flow [15, 18–20], and the energetic barriers for T1 transitions to occur can dictate tissue fluidity/solidity [9, 21–23].

For a cell monolayer under 3D geometric constraint, cells can undergo apical-basal T1 (AB-T1) transitions (Fig. 1A, top). Different from the planar and dynamic T1-transitions described above, AB-T1 transitions are a static exchange of neighbors from the apical to basal layers of the cell. Such a 3D cellular arrangement, termed as a *scutoid* in the context of epithelial tissues [24–26] (Fig. 1A), has been observed in foams [27, 28] and biological systems with curved surfaces [29–33].

**FIG. 1.**
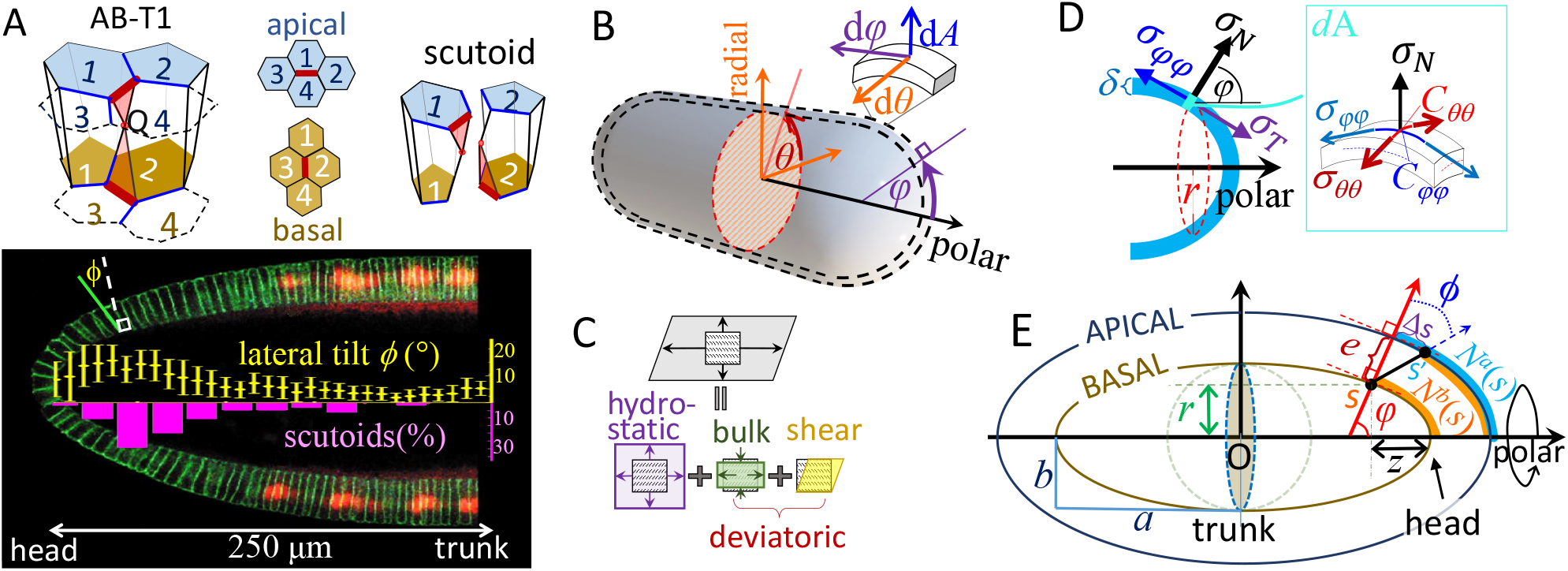
The effect of curvature on cell packing and cellular forces. (A) Top: Scutoid geometry in epithelial tissues; point *Q* is the additional point shared by two columnar cells. The AB-T1 transition occurs at the edges highlighted in thick brown, leading to the exchange of neighbor pair from 1-4 (apical) to 2-3 (basal). Bottom: Tilt angle of lateral membrane (yellow) and percentage of scutoids (pink) peak near the head of a wild type *Drosophila* embryo, adapted from [29] under a Creative Commons License. (B) Two-layered model for curved epithelia on an axisymmetric object and the coordinates for a any local surface *dA*; (C) Graphical representation of the stress tensor decomposition, Eq. (1). (D) Force balance of a curved layer under axisymmetric loads: (left) at the meridional cut (red dashed ring) and (right) along the normal direction of the element surface *dA*(*φ, θ*). (E) A meridional cross section view of a two-layered prolate ellipse. The black tilted line is the tilted lateral membrane, with the basal end at *s* and the apical end at *st*, with the tilt angle *ϕ* and apical-basal distance *e* at *s*. The orange curves are the accumulated cell number from the head to *s* at the basal side; the skyblue curve is the accumulated cell number from the head to *s* at the apical side.

Tissue curvature is proposed to be pivotal in inducing AB-T1 transitions. In the ellipsoidal early *Drosophila* embryo, AB-T1 transitions appear most frequently around 20-50*μ*m from the embryo head, a region with low curvature anisotropy but large tilt of cell lateral membranes [29] (Fig. 1A). During salivary gland formation in the *Drosophila* embryo, AB-T1 transitions occur at maximal curvature anisotropy [24]. Models have been proposed for cell packing in these specific cases [24, 29], but there is currently no consensus on how curvature induces AB-T1 transitions.

Here, we provide a framework for describing curvature-induced cell deformation, which can be generalized to an array of geometries, and discuss the interplay between cell mechanics and tissue geometry in inducing AB-T1 transitions. We demonstrate that in 3D environments with steep curvature gradient, cells can tilt in order to pack efficiently. These tilted lateral membranes can exert tensions that contribute to in-plane stresses of opposite sign on the apical and basal plane stresses, thereby leading to AB-T1 transitions. Conversely, when hydrostatic pressure dominates, we find that AB-T1 transitions occur in regions with high curvature anisotropy. Overall, we find that the combination of tissue curvature, pressure, and lateral tensions determines the location of AB-T1 transition events.

## Framework

We treat the epithelia as a material composed of two connected thin shells, representing the apical and basal surfaces of the tissue. Assuming the radius of curvature to be significantly larger than the cell size, we can use a continuum mechanics model based on membrane theory for elastic thin shells, neglecting bending stresses. Lateral membranes are included as part of the external load on the shell. Motivated by the *Drosophila* embryo, salivary gland and oocyte geometries, we focus on axisymmetric geometries, which have rotational symmetry about a polar axis (Fig. 1B). For any infinitesimal surface element *dA* on the 3D curved shell, it has a normal direction *dA*, and two tangential directions along the meridian *dφ* and latitudinal radii *dθ* (Fig. 1B).

The in-plane stresses in the apical or basal layer are described as a stress tensor 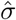 bearing two principal stresses *σ*_*φφ*_, *σ*_*θθ*_ and a shear stress component 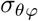, with the basis 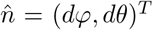. This stress tensor 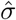 can be decomposed into a hydrostatic part 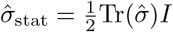, corresponding to isotropic forces that induce local expansion or shrinkage of cell areas, and a deviatoric part 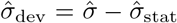 corresponding to the anisotropic forces that induce shearing or anisotropic bulk compression/stretching (Fig. 1C).

The above stresses are balanced by the external loads from the lateral and apical/basal membrane generated by cell deformation or cellular active forces [34, 35]. For simplicity, we only consider axisymmetric external load, which can be decomposed into a normal part *σ*_*N*_ (positive pointing outward) and a tangential part along the meridian *σ*_*T*_ (positive pointing to the head) and hence the in-plane shear 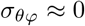. The meridional stress *σ*_*φφ*_ at any local cut (red ring in Fig. 1D) is balanced in the polar direction by the accumulated force over the revolved surface as:

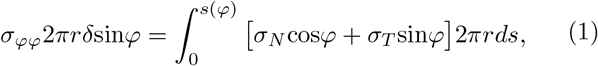

where *δ* is the thickness of cell membrane, *r* is the distance to the polar axis from the local surface *dA* (Fig. 1D) and *ds* is the meridional arc length (for derivation, see Supp. Mat. A). The circumferential stress *σ*_*θθ*_ is derived from force balance along the normal direction of the surface:

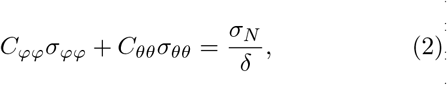

where *C*_*φφ*_ and *C*_*θθ*_ are the principal curvatures along the meridional and circumferential direction, respectively.

### AB-T1 transitions

The stresses in apical or basal layers can induce cell shape changes and cell intercalations. Here, we assume that prior to any applied external load, cells are relaxed to isotropic shapes without any deviatoric strain. AB-T1 transitions will take place most frequently when the apical and basal sides of a cell have oppositely directed deviatoric stresses [36] under external loading. In the absence of shear components *σ*_*φθ*_, we can define a measure for AB-T1 transitions, *γ*, as proportional to the difference of the deviatoric strain between the apical and basal sides:

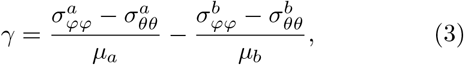

where *μ*_*a,b*_ represent the effective elastic moduli at the apical and basal surfaces; *γ >* 0 corresponds to cells that are stretched along the meridional direction at the apical side while compressed along the circumferential direction at the basal side. The parameter-dependence of *μ*_*a,b*_ depends on the underlying material properties. As demonstrated in Supp. Mat. B, taking different forms for *μ*_*a,b*_ does not alter our key conclusions. Here, we consider 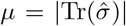, which avoids introducing an intrinsic elastic modulus for the cells. Under typical physiological regimes for epithelial cells, we expect 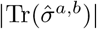 to be non-zero, so *γ* behaves well.

We first consider the case when external loads are hydrostatic (*σ*_*T*_ = 0 and *σ*_*N*_ = *P*). With large curvature anisotropy, |*C*_*θθ*_ − *C*_*φφ*_|, the magniture of *γ* is large, leading to AB-T1 transitions. In contrast, isotropic curvatures (*C*_*θθ*_ = *C*_*φφ*_) lead to *γ* = 0 (derivations in Supp. Mat. C). This conclusion is consistent with the experimental observations in tubular epithelia [24].

### Cell tilting

The results for hydrostatic systems above are not consistent with the AB-T1 transitions observed in the head of the early *Drosophila* embryo [29], where the curvature is nearly isotropic. However, in this system, the cells are observed to tilt (Fig. 1A). The profile of external load *σ*_*T*_, *σ*_*N*_ is affected by tilt of lateral membranes. We next investigated cell tilting within our model and explain its role in inducing AB-T1 transitions. The tilted lateral membrane leans to the head by a small angle *ϕ* away from the normal direction (illustrated in Fig. 1E) as

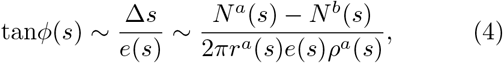

where Δ*s* is the distance between the apical projection of *s* and the apical end of the tilted lateral membrane *s*′; *e*(*s*) is the distance between the apical and basal layer; *ρ*^*a,b*^(*s*) is the cell density; *r*^*a,b*^(*s*) is the distance from *s* to the polar axis; 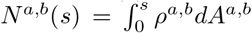are the accumulated number of cells from the head apex to the coordinate *s* on the apical and basal sides, respectively. Although Fig. 1E is illustrated for an prolate ellipsoid, Eq. 4 works for any arbitrary axisymmetric shape.

The distribution of *ρ*^*a,b*^(*s*) and *e*(*s*) are interdependent, as a consequence of minimizing the system free energy including the contributions from cell lateral membranes (Supp. Mat. E). If the lateral membrane tensions are weak compared with the apical and basal cell layers, the apico-to-basal density ratio *ρ*^*a*^(*s*)*/ρ*^*b*^(*s*) converges to a space-independent constant (Supp. Mat. D). In this limit, the tilt angle

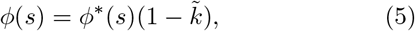

where 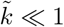 is the ratio of tension strength between the lateral and apical/basal layers; *ϕ*^*\**^ is the tilt in the limit of zero lateral tension, depending on the curvature as:

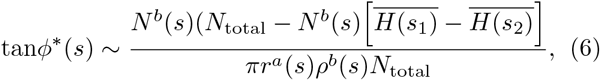

where 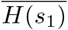 and 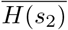are the mean curvature weighted by cell numbers in a range of 0 *< s*_1_ *< s* and *s < s*_2_ *< s*_0_, respectively (*s*_0_ is the half meridian). For a convex object, a large gradient of *H*(*s*) corresponds to a large magnitude of *ϕ*^*\**^ at *s*, with the corresponding tilt direction towards the region of higher positive curvature (Supp. Mat. E).

Conversely, if lateral membranes are extremely rigid, the lateral membrane tends to stand perpendicular to the surfaces, and *ρ*^*a*^(*s*)*/ρ*^*b*^(*s*) equals inverse apico-to-basal area ratio *dA*^*b*^(*s*)*/dA*^*a*^(*s*), hence the tilt vanishes (Supp. Mat. G). To further simply the model, we show that the effect of any cell density inhomogeneity on cell tilt is negligible if cell density changes along the surface more slowly than the curvature does (Supp. Mat. E). We henceforth set a homogeneous density 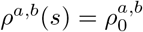.

### Ellipsoid case

We now apply this formalism to a prolate ellipsoidal geometry as shown in Fig. 1E. It has a major half axis *a* and minor half axis *b* (see Supp. Mat. F for parameterization and the calculation of the curvature). Tissue height is determined mainly by the intrinsic cell volume control [37]. To leading order in the arc length *s* to the head, the height profile reads

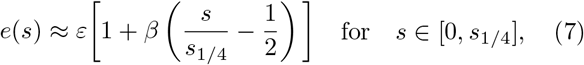

where *s*_1*/*4_ is the 1*/*4 perimeter of the meridian ellipse and *ε* is the average cell height across the surface and *β* is a coefficient modulating the surface height with *β* = 0 representing homogeneous cell height. As we assume cell size is much smaller than the radius of curvature, the average height of the tissue *ε* has negligible impact on the tilt profile (see discussions in Supp. Mat. E).

We calculate the cell tilt angle *ϕ*^*\**^ in the zero-lateral-tension limit as a function of the relative distance to the head of a prolate ellipsoid, *d*(*s*) = *z*(*s*)*/a*, where *z*(*s*) is the distance to the head along the polar direction; *d* = 0 corresponds to the head and *d* = 1 to the trunk. The tilt angle increases with elongation of the ellipsoid (smaller *b/a*), Fig. 2A. For a typical value observed experimentally in *Drosophila* (*b/a* ∼ 0.4 [29]), the tilt angle peaks around 30°. The impact of height inhomogeneity on the tilt angle is shown by Fig. 2B: a large, positive *β* (tissue height larger at the trunk) makes the peak of the tilt angle profile more pronounced. The calculated tilt profile is consistent with the data observed in the early *Drosophila* embryo (*β* ∼ 0.5), with the predicted magnitude of *ϕ*^*\**^ (red curve) slightly larger than the experimental measurements (black dots, from [29]) as expected by Eq. 5.

**FIG. 2.**
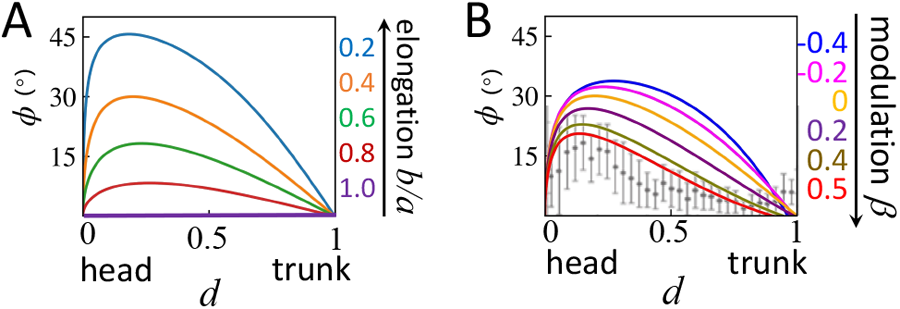
The lateral tilt angle in the zero-lateral-tension limit, Eq. 6, as a function of the distance to the head along the polar direction *d* = *z/a* (Fig. 1E) (A) under varying inverse aspect ratio *b/a* at *ε/a* = 0.05, *β* = 0; (B) under varying thickness modulation *β* at *ε/a* = 0.01, *b/a* = 0.4. Experimental data is shown for the cell tilt angle in the early *Drosophila* embryo (*b/a* ∼ 0.4, *β* ∼ 0.5), with s.d., by grey dots (data from [29]).

External loads along the tilted lateral membranes can qualitatively change the stress distribution. We show in Fig. 3A-B a comparison of the stress components *σ*_*φφ*_ and *σ*_*θθ*_ between a hydrostatic case: 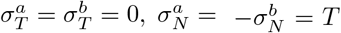 and a case with the external stresses *T* along tilted lateral membranes:

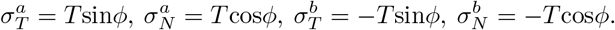

**FIG. 3.**
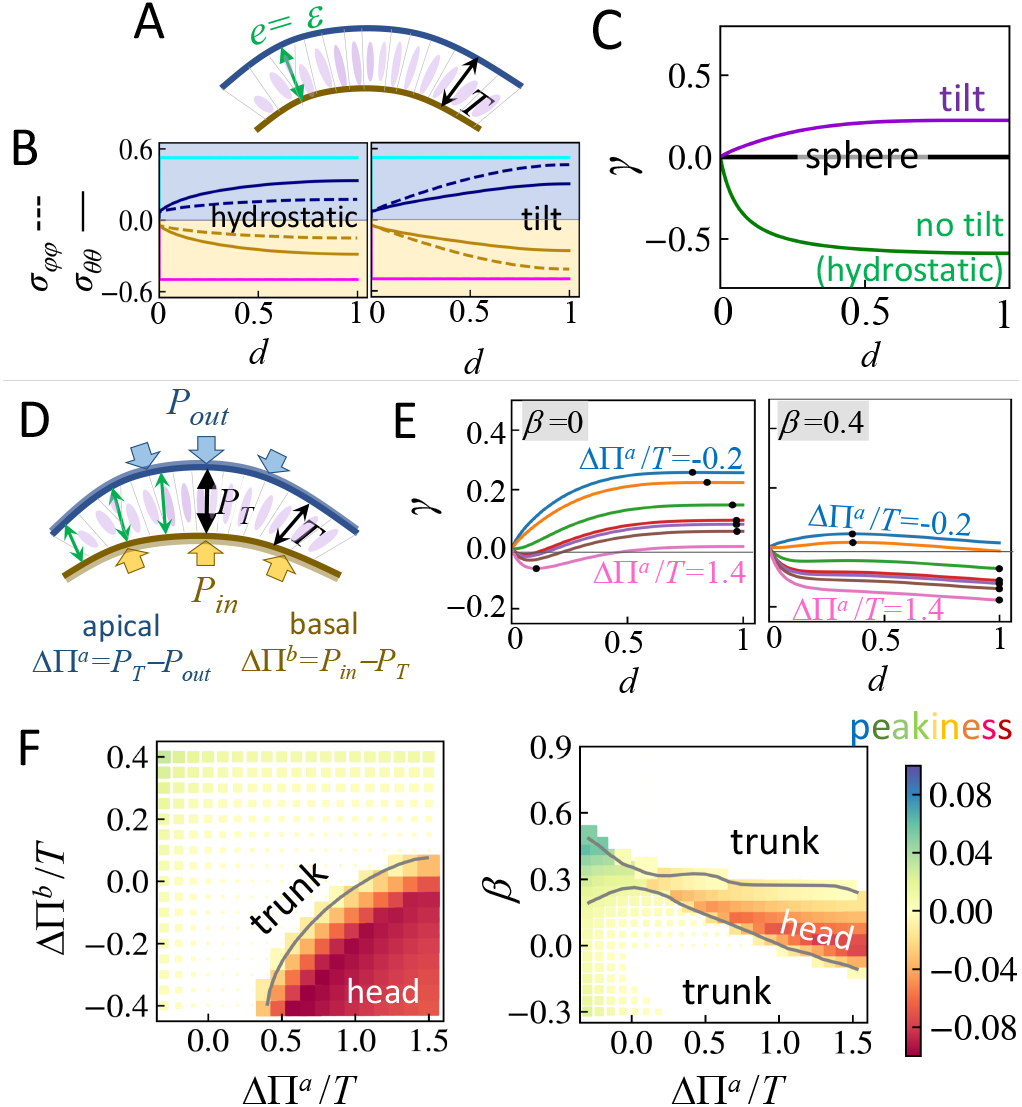
AB-T1 transition rate *γ* calculated for a prolate ellipsoid with *b/a* = 0.4. (A) A schematic illustration of the system under tensile lateral stress *T* with a constant tissue height *ε* = 0.05*a* for panel B and C; (B) In plane, apical and basal stress components normalized by *Ta/δ* as a function of the distance to the head. Left: *T* perpendicular to the layer (hydrostatic); right: *T* along tilted lateral membranes. The cyan and magenta curves stand for a sphere (*a* = *b*). The tensile stresses have a positive sign (apical, blue) and contractile stress has a negative sign (basal, yellow). (C) The correspondent AB-T1 rate *γ* for the prolate ellipsoid. (D) A schematic tissue setting under the external loads with pressures and inhomogeneous tissue height (Eq. 7). (E) Profile of the AB-T1 measure, *γ*, with varying apical pressure difference ΔΠ^*a*^ and with basal pressure difference ΔΠ^*b*^ = 0. Left: the tissue height modulation rate *β* = 0; right: *β* = 0.4. Black dots indicate the peak, where the absolute AB-T1 measure |*γ*| reaches the maximum. (F) The phase diagram for the peakiness of *γ*, which is calculated as sign(*γ*_peak_) × ||*γ*_peak_|− |*γ*_trunk_||. Left: *β* = 0; right: ΔΠ^*b*^ = 0. The size of the scattered square is ∝ (1− *d*_peak_) ^2^, so positions closer to the head (*d* = 0) are represented by larger squares. The grey contours separate the trunk (*d*_peak_ *>* 0.5) and the head (*d*_peak_ *<* 0.5) regions.

The magnitude of *σ*_*θθ*_ and *σ*_*φφ*_ grows from the head to the trunk in different manners, depending on whether *T* is perpendicular to the shells (hydrostatic) or *T* along the tilted lateral membranes. The resultant AB-T1 transition rate, calculated through Eq. 3, flips its sign with or without the tilt (Fig. 3C). However, this qualitative difference will vanish when the surface approaches a sphere (*a/b* = 1) (Fig. 3B; cyan and magenta lines), leading to no AB-T1 transition at all locations (Fig. 3C; black line). Next, we discuss results with pressure differences across the cell along the apical-basal axis (Fig. 3D). The tissue height follows Eq. 7. The apical and basal membranes are subject to pressure from: the outside *P*_*out*_; from the internal cavity (*e*.*g*. yolk or luminal pressure) *P*_*in*_; and inside the tissue *P*_*T*_. The pressure differences at the apical and basal surfaces are given by ΔΠ^*a*^ = *P*_*T*_ −*P*_*out*_ and ΔΠ^*b*^ = *P*_*in*_ −*P*_*T*_ respectively, with positive ΔΠ pointing towards the outside. Before applying external load, we assume cells have relaxed to their preferred cell shape with no internal strain. The external normal and tangential loads on the apical and basal side are 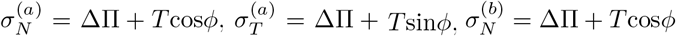 and 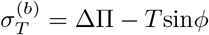.

The system dominated by pressure (ΔΠ*/T*→∞) corresponds to a hydrostatic limit, Fig. 3B (left). In this limit, the profiles of stresses and the consequent spatial distribution of AB-T1 transition frequency do not qualitatively depend on the pressure differences or the cell height profile (Supp. Mat. C). In contrast, strikingly, when the pressure difference is comparable with lateral stress (ΔΠ ∼*T*), *γ* is sensitive to the two pressure differences and *β*, Fig. 3E. ΔΠ can be negative (pointing inwards), thus the normal component of lateral tension *T* can be partly balanced by this pressure and *σ*_*T*_ */σ*_*N*_ becomes much larger as if the cells tilt more significantly. When ΔΠ^*a*^*/T* shifts sign from negative to positive, at the trunk (*d* = 1) *γ* becomes negative, altering the orientation of AB-T1 transitions.

To capture the key features of the distribution of |*γ*|, we define the peak of *γ* as where *γ*_peak_ is the maximal value of *γ* (Fig. 3E) and its value at peak (referred here as the *peakiness*) as sign(*γ*_peak_) ×||*γ*_peak_| −|*γ*_trunk_||. Accordingly, we can construct a phase diagram of AB-T1 transitions, using the position of the peak and peakiness as the order parameters, Fig. 3F. We show the diagram in the ΔΠ^*a*^-ΔΠ^*b*^ space for *β* = 0 (left) and in the space of *β*-ΔΠ^*a*^*/T* with ΔΠ^*b*^ = 0 (right). The peak in the tendency of AB-T1 transition switches from trunk to head beyond a critical line *β*(ΔΠ) (Fig. 3F). From these phase diagrams, we can estimate mechanical properties (*e*.*g*. pressure, lateral tension, or possible external loads) from the geometric cell profiles (*e*.*g*. cell tilt, cell height and AB-T1 locations/orientations).

## Conclusions

We have proposed a model for the onset of cellular tilt within a curved monolayer. We find that the interplay between the lateral cell-cell tension and the cellular tilt leads to a shift in the location at which we expect the number of neighbor rearrangements to be maximal. Our formalism provides predictions for the location of AB-T1 transitions in several geometries that are echoed by experimental observations in various geometries [24, 29].

The lateral membranes play an essential role in balancing stress across the cell, thereby regulating cell shape. In particular, lateral membranes with low contractility lead to cell tilting, which cooperates with pressure and tissue thickness to result in a rich phase diagram for the tendency of AB-T1 transitions to occur. If the lateral membranes are sufficiently stiff, then the tilt of lateral membranes is suppressed and AB-T1 transitions occur at regions with large curvature anisotropy, following the model prediction in the hydrostatic limit.

Though we have focused on a prolate geometry with simple external loads, our formalism can be generalized to a diverse range of tissue geometries observed *in vivo*. We expect tilt to occur at the steepest curvature gradient, even for non-axisymmetric and non-closed surface geometries; *e*.*g*. the brain and gut. We can also explore the role of in-plane shear and bending within this theoretical framework. Internal cell strain, which is likely significant during cellular process such as cell division[38], can also be considered as a source of external loading. Finally, transient and reversible AB-T1 transitions have been observed [39, 40]; the dynamic aspect of AB-T1 transitions may be relevant to the mechanism of T1 transitions [23, 36] and their contributions to processes like tissue folding or buckling [41−46] remains to be investigated.

We thank Jacques Prost for discussion leading to Eq. (4). J.-F.R. is funded by ANR-16-CONV-0001 and ANR-20-CE30-0023 grants and from the Excellence Initiative of Aix-Marseille University - A*MIDEX. T.H. and T.E.S. are funded by Mechanobiology Institute seed grants. T.E.S. is also funded by start-up support from the University of Warwick.

## SUPPLEMENTARY MATERIAL

### A. Force balance in axisymmetric systems

For a general elastic material, the force balance in terms of the stress tensor 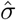 is

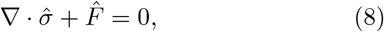

where 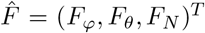is the external body force exerted on a element of a small volume. For a thin elastic shell revolved around a polar axis with distance *r*(*s, θ*), this body element = *dA · δ*, where *dA* = *rdsdθ* and *δ* represents the thickness. *ds* is an infinitesimal length along the meridional direction and *rdθ* is an infinitesimal length along the circumferential direction. The radius of curvature along *ds* and *rdθ* are denoted as *R*_*φφ*_ and *R*_*θθ*_, respectively. Hence *ds* = *R*_*φφ*_*dφ* and *R*_*θθ*_ = *r/*sin*φ*.

For a thin shell (*i*.*e. δ* is much smaller than the typical curvature radius of the system), these quantities can be taken as uniform along the thickness direction, so the transverse shear can be neglected. Therefore, only the inplane stresses *σ*_*φφ*_, *σ*_*θθ*_ and *σ*_*φθ*_ are considered in the force balance and their derivatives along the normal directions are neglected. Furthermore, we do not consider possible bending stresses at the discontinuity of displacement (usually at the apex of the object) due to the in-plane stresses [47–49]. With these assumptions, Eq. 8 leads to a set of force balance equations along the meridional 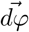, circumferential 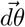and normal directions to the surface 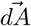:

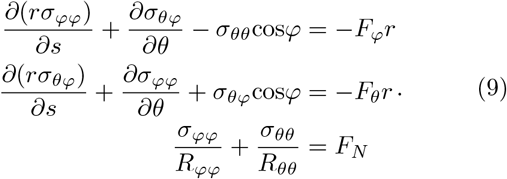

For an axisymmetric system *r*(*φ, θ*) = *r*(*φ*), we drop all the terms with derivatives with *∂θ* and obtain the axisymmetric resultant for the top and bottom equations in Eq. 9 as

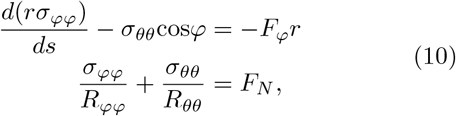

which are independent of shear. The torsion around the polar axis from shear force is exclusively determined by the second equation in Eq. 9. For our case, *F*_*θ*_ = 0 so the shear component must also be zero throughout the space considering the boundary condition *σ*_*θφ*_(*φ* = 0, *π*) = 0.

From Eq. 10, we obtain a differential equation for *σ*_*φφ*_

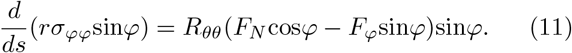

Integrating and multiplying Eq. 11 by a factor 2*πδ*:

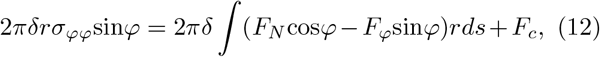

where the left hand-side is the total force parallel to the polar axis found at a latitudinal cross-section of the shell positioned with arc length *s*. This force is balanced by the distributed load across the surface along with a concentrated force *F*_*c*_ at the apex *s* = 0. The indefinite integral could be alternatively expressed by a definite integral from *s* = 0 to *s* = *s*(*φ*).

Here, there is no reason to consider a concentrated force at the head apex, so we set *F*_*c*_ = 0. We define the loading from the two external stresses *σ*_*T,N*_ per unit area such that *δF*_*φ*_ = −*σ*_*T*_ (with a minus sign so that *σ*_*T*_ *>* 0 points towards the head) and *δF*_*N*_ = *σ*_*N*_. With these notations, Eq. 12 is equivalent to Eq. 1 in the main text.

### B. Measure for AB-T1 transition likelihood

We define a measure *γ* for the tendency of finding a AB-T1 transition as the difference in the deviatoric strain between apical and basal layers (Eq. 3). The magnitude of *γ* relates to the probability of finding an AB-T1 transition, and the sign of *γ* indicates the orientation of the corresponding AB-T1 transition, as described in the text. The deviatoric strain is the deviatoric stress [36, 50], divided by an effective tissue shear modulus *μ* as:

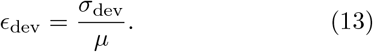

Here, we ignore shear and torsion so *σ*_dev_ = *σ*_*φφ*_−*σ*_*θθ*_. A positive sign indicates a tensile strain along the meridian direction with a compressive strain along the circumferential direction.

The effective shear modulus *μ* is related to the strength of the tissue in resisting deformation in exchanging neighbors along the AB direction. This modulus depends on how the cell cortex biopolymers connect, bend, and interact in the material. Some empirical and theoretical literature has shown that the shear modulus of tissues is stiffened by pre-compression or pre-expansion of the tissue [51–53]. Tension stiffening originates from a bending-to-stretching mode transition, while the mechanism of compression stiffening originates from jamming [53, 54]. In some particular cases, a tissue can even display tension strain-softening due to connections breaking between adherent regions [54].

Supposing that the pre-stress in plane is small, we have a phenomenological linear relationship for the effective shear modulus

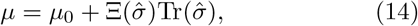

where *μ*_0_ is the intrinsic shear modulus of the material and the trace of in plane stress tensor indicates the isotropic tensile or compressive stresses in the layer. The dimensionless coefficient Ξ can have various dependencies on the stresses 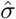 for a broad range of hyperelastic materials. Here, we discuss several simple forms for Ξ.

**FIG. S1.**
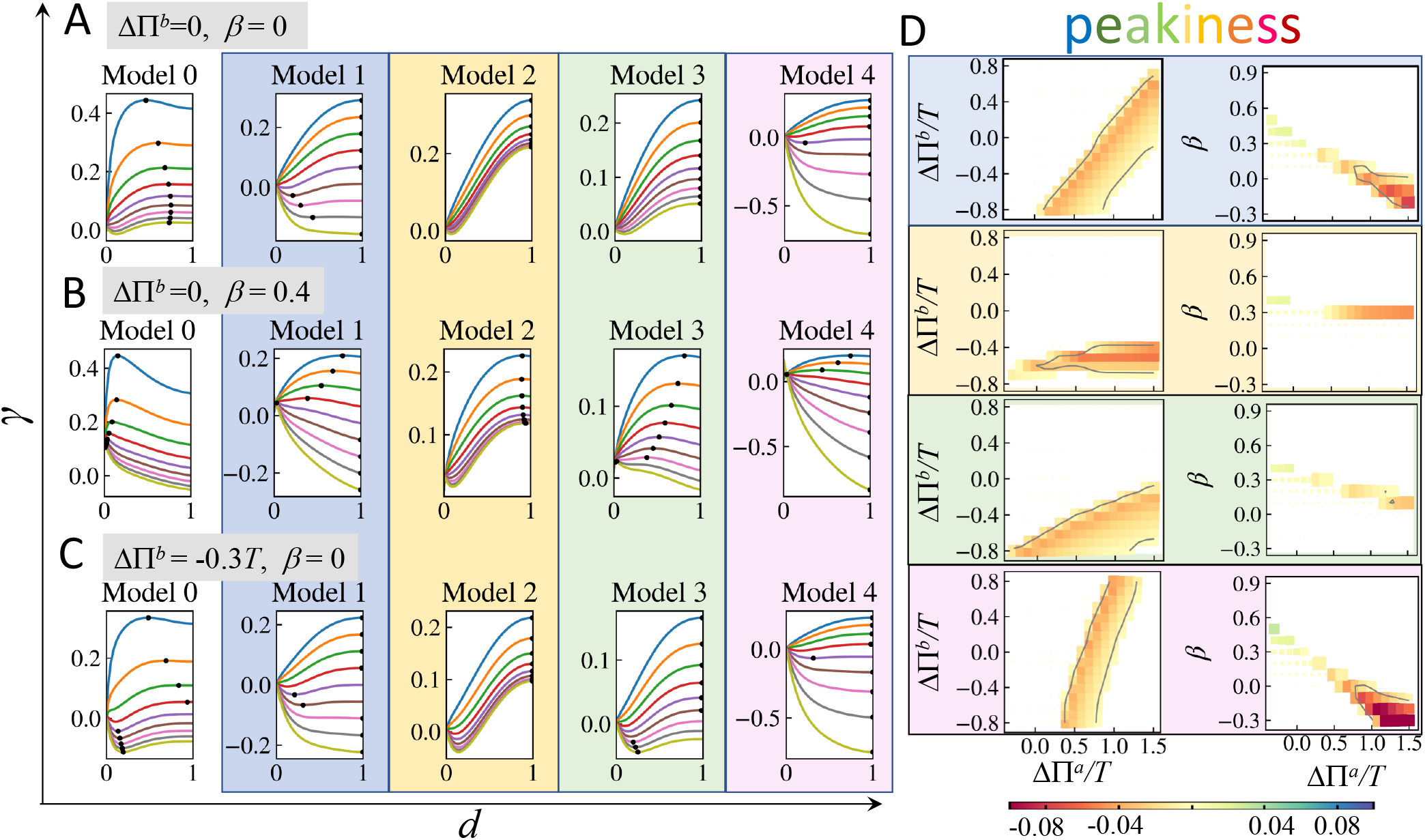
Comparison among different models for effective shear modulus for a prolate ellipsoid *b/a* = 0.4. (A-C) Distribution of *γ* for varying apical pressure differences ΔΠ^*a*^*/T* from top to bottom: −0.3 (skyblue), 0 (orange), 0.3 (green), 0.6 (red), 0.9 (purple), 1.2 (brown), 1.5 (pink), 1.8 (grey), 2.1 (golden), with different values of the basal pressure difference ΔΠ^*b*^ and tissue height inhomogeneity *β*. Horizontal axis *d* is the relative distance; *d* = 0 indicates the head of a prolate ellipsoid and *d* = 1 the trunk. Black dots indicate the peak of |*γ*|. (D) Diagrams of peakiness, which is defined as sign(*γ*_peak_) × ||*γ*_peak_| − |*γ*_trunk_||. From top to bottom: Model 1, Model 2, Model 3, Model 4. White regions indicate peak of |*γ*| at trunk, whereas coloured squares indicate peak of |*γ*| at near the head. The corresponding diagrams for Model 0 are presented in the main text Fig.3F.

First of all, we consider a linear prestress-stiffening, modeled by a positive constant *ξ* for tensile stresses (trace of stress tensor *>* 0) and a negative constant −*ξ* for compressive stresses (trace of stress tensor *<* 0); hence Eq. 14 becomes

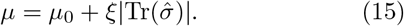

By tuning the value of *ξ*, one can explore varying effects of prestress-stiffening in the model.

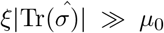 (a strong prestress-stiffening), the intrinsic shear modulus can be ignored such that

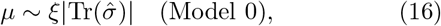

which, normalized by *ξ*, is used for the results shown in the main text.

Oppositely, if 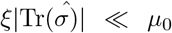(negligible prestress-stiffening), the effective shear modulus is dominated by the intrinsic shear modulus such that

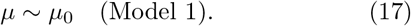

There are other simple forms for *μ*: (i) tension-stiffening while compression-softening:

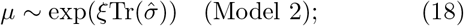

(ii) only tension-stiffening;

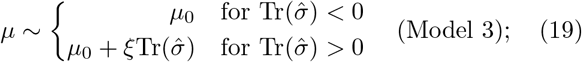

or (iii) only compression-stiffening:

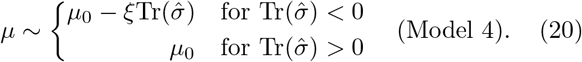

The comparison of *γ* under these five types of effective shear modulus are shown in Fig.S1. External load acts through pressures differences ΔΠ^*a*^ and ΔΠ^*b*^ at apical and basal sides, along with the lateral tension *T*, as demonstrated in the main text (Fig. 3D). In Models 2, 3, 4, we set *μ*_0_ = *ξ* and all the *γ* shown here are normalized by *a/δξ*. In Fig.S1D, we show the diagrams of peakiness as defined in the main text in the parameter space of ΔΠ^*b*^*/T* − ΔΠ^*a*^*/T* and *β* − ΔΠ^*a*^*/T* for Models 2-4.

We can see for different models of the effective modulus *μ*, the phase diagrams of peakiness have different boundaries between the trunk region (white) and the head region (colored) in the parameter space. Both the material properties and the tissue geometry play important roles in the occurrence and positioning of the AB-T1 transitions. To distinguish the models, we refer back to the experimental observations. In the early *Drosophila* embryo, AB-T1 transitions are very infrequent at the anterior head of the embryo and also in the trunk region. Comparing the *γ* distribution between the different models, we see that a distribution with peak in *γ* near (but not at) the head with near zero value in the trunk (*γ*(*d* = 1)*/γ*_peak_ ∼0) has a very narrow parameter space in all the models. This is because *γ* ∼0 at the trunk requires pressure and stresses along the lateral membrane to be closely balanced.

### C. Deviatoric strain under hydrostatic load

In this section, we show the analysis of the deviatoric strain profile and the correspondent AB-T1 likelihood under the hydrostatic external loads. Substituting *σ*_*T*_ = 0, *σ*_*N*_ = *P* into the force balance equations Eqs.1-2, we arrive at:

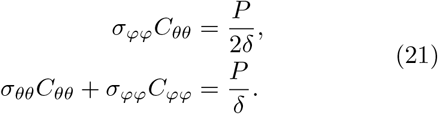

Solving Eq. 21 yields the following relations:

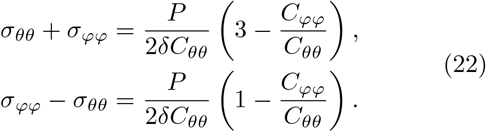

Substituting Eq. 22 into a general expression of the deviatoric strain, with the *μ* defined as in Eq. 15 in Supp. Mat. B, we obtain the following analytical expression of the deviatoric strain for a layer:

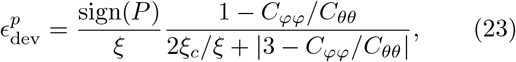

where *ξ*_*c*_ = *δC*_*θθ*_*μ*_0_*/*|*P* |.

Note that for an axisymmetric system, *C*_*θθ*_ is always positive, *i*.*e*. the small arc along the circumference is always convex to the polar axis, while *C*_*φφ*_ can either be positive for a convex meridian or negative for a concave one, with respect to the polar axis. Figure S2 is a graphical representation of a normalized deviatoric strain 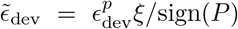 against *C*_*φφ*_*/C*_*θθ*_. When *C*_*φφ*_*/C*_*θθ*_ = 1, the two prime curvatures of a local surface are the same and under hydrostatic external loading, 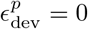, independent of *ξ*.

**FIG. S2.**
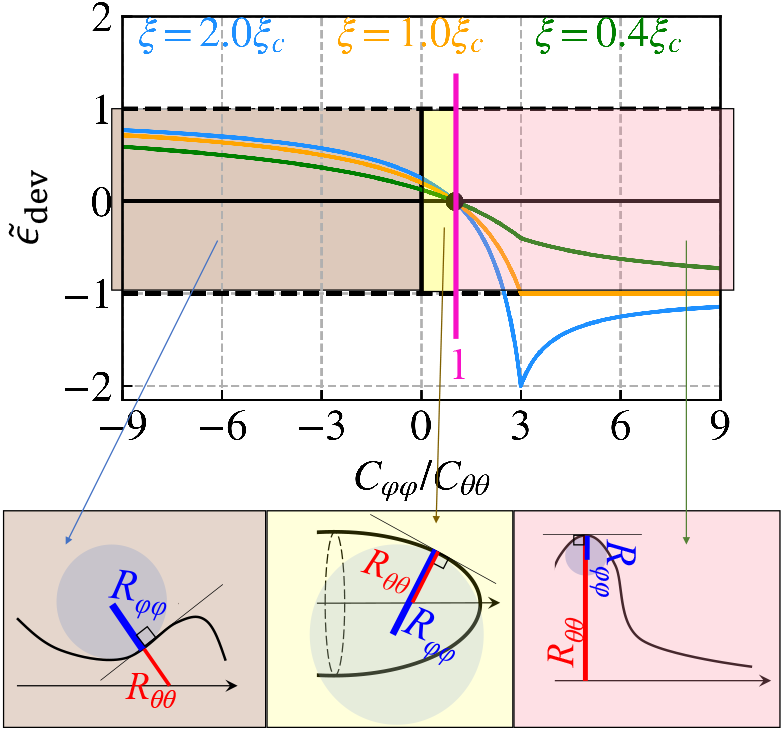
The normalized deviatoric strain 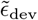 in relation to the ratio of prime curvatures *C*_*φφ*_*/C*_*θθ*_. All 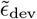 with various *ξ* at isotropic curvature condition *C*_*φφ*_ = *C*_*θθ*_ (magenta line). There are three typical examples of different curvature ratio below: brown (left) for *C*_*φφ*_*/C*_*θθ*_ *<* 0; yellow (middle) for 0 *< C*_*φφ*_*/C*_*θθ*_ *<* 1; pink (right) for *C*_*φφ*_*/C*_*θθ*_ *>* 1. The arrow indicates the polar axis. The radius of curvature *R*_*θθ*_ = 1*/C*_*θθ*_ and *R*_*φφ*_ = 1*/C*_*φφ*_ are highlighted by the red and blue lines respectively.

When *ξ* ≫ *ξ*_*c*_ (strong stress-stiffening, blue curves in Fig. S2), the largest magnitude of 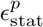 occurs at *C*_*φφ*_*/C*_*θθ*_ = 3 with its value

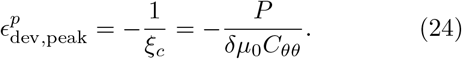

For a shape elongated along the polar axis without bumps, |*C*_*φφ*_ − *C*_*θθ*_| = 3 is not feasible. In this case, the magnitude of *E* increases with *C*_*φφ*_*/C*_*θθ*_ → −∞, where the curvature anisotropy |*C*_*φφ*_ − *C*_*θθ*_| becomes large.

When *ξ* « *ξ*_*c*_ (weak stress-stiffening, green curves in Fig. S2), the magnitude of 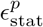 always increases with the growth anisotropy of curvature. The largest magnitude of deviatoric strain occurs with value 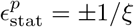 when *C*_*φφ*_*/C*_*θθ*_ → ∓∞.

When *ξ* ∼ *ξ*_*c*_ = *δμ*_0_*C*_*θθ*_*/*|*P* | (orange curve in Fig. S2), the largest magnitude of 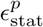 occurs where *C*_*φφ*_*/C*_*θθ*_ *>* 3. Large *C*_*φφ*_*/C*_*θθ*_ corresponds to geometries such as bumps, see the right bottom panel in Fig. S2.

Although the strength of prestress-stiffening (value of *ξ*) affects the magnitude of deviatoric strain in different ways for *C*_*φφ*_*/C*_*θθ*_ *>* 1, the behaviors of deviatoric strain are robust against *ξ* for the region *C*_*φφ*_*/C*_*θθ*_ *<* 1, which is typically the curvature ratio for a regular elongated axisymmetric shape, such as an ellipsoid and cylindrical tube. In these systems, the largest magnitude of deviatoric strain occurs at the smallest value of *C*_*φφ*_*/C*_*θθ*_. If we narrow the cases to only convex surfaces, *i*.*e. C*_*φφ*_ *>* 0 (see the middle bottom panel in Fig. S2), then the largest magnitude of deviatoric strain occurs at *C*_*φφ*_ = 0. This corresponds to the trunk region of a prolate ellipsoid. This conclusion also holds when considering the other forms of the effective shear modulus *μ*^*a,b*^ (*e*.*g*. those discussed in Supp. Mat. B).

Let *c* denote the ratio between two principal curvatures *C*_*φφ*_*/C*_*θθ*_. Under hydrostatic situation, the AB-T1 tendency profile *γ*, which is the difference of deviatoric strain between apical and basal side, becomes

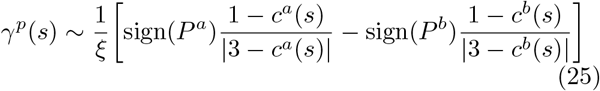

for *ξ/ξ*_*c*_ ≫ 1 and

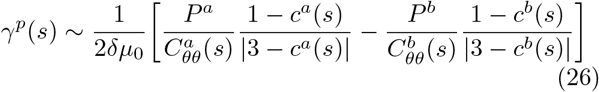

for *ξ/ξ*_*c*_ ≪ 1.

Since the cell height *e* is much smaller than the radius of curvature (our model assumption), 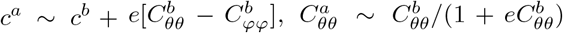, then the profile of *γ*^*p*^(*s*) is approximately proportional to the normalized hydrodtatic deviatoric strain at the basal side 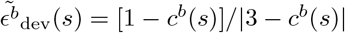 as

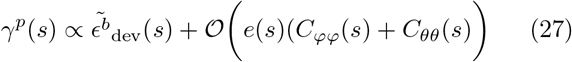

with a linear coefficients determined by the hydrostatic loading *P* ^*a,b*^ and a negligibly small correction from the cell height. Hence, the AB-T1 tendency under hydrostatic conditions, *γ*^*p*^, is near zero at locations with isotropic curvature and increases with the curvature anisotropy.

### D. Calculation of tilt angle of the lateral membrane on an arbitrary axisymmetric object

Fig. S3 illustrates a meridional cross section for an arbitrary axisymmetic shell. A surface element *dA*(*φ, θ*) located at *s* on the basal side (golden in Fig. S3), can be mapped to another surface element *dA*^*a*^(*φ*′, *θ*′) located at *s*′ in such a way that the accumulated number of cells from the head of the object to *s* on the basal side is the same as the cell number accumulated from the head to *s*′ at the apical side. Hence, the angle *ϕ* between the normal direction of the surface and 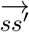 is the tilt angle of the cells at *dA*. The surface element *dA* = *r*(*s*)*dθds*, where *r*(*s*) is the distance to the polar axis, and *rdθ* and *ds* are the two orthogonal vectors along the circumferential and meridional directions, respectively.

Given our assumption of an axisymmetric surface, the 2D integral of surface element *dA* over the whole shell surface can be reduced to a 1D integral with only the meridional variable from 0 to *s*. The accumulated number of cells *N* from the head to *s* on the basal side is given by

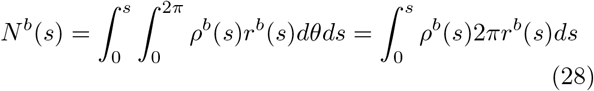

and it is equal to the accumulated number of cells *N*^*a*^ on the apical side:

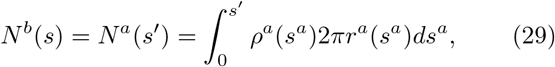

where *ρ*^*b*^(*s*) or *ρ*^*a*^(*s*) is the cell density on the basal or apical surface, *r*^*b*^(*s*) or *r*^*a*^(*s*) is the circumferential radius at *s*. The density *ρ*^*a,b*^(*s*) is determined by minimizing the membrane tensions on apical, basal and lateral sides. Here, we do not consider any other cues guiding cell location within the tissue environment.

**FIG. S3.**
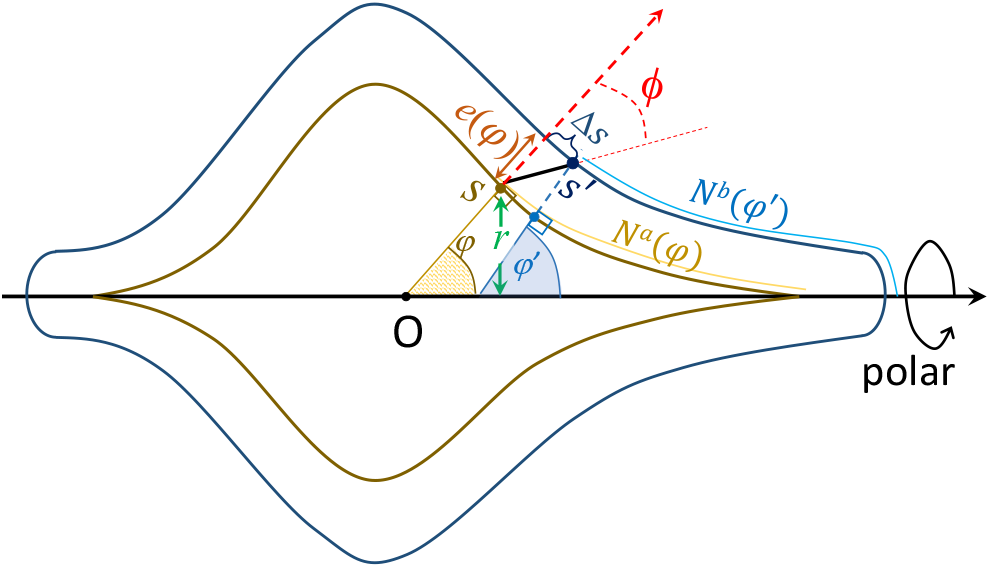
A meridional cross section of an arbitrary axisymmetric shell with apical (navy) and basal (golden) layers. The apical layer is an outward projection of the basal layer along the normal direction at each local surface element with a distance *e*. The position *s*′on the apical side corresponds to position *s*′ on the basal side in such a way that the cell number accumulated on the apical surface from the head to *s*′ equals the basal one accumulated from head to *s*; therefore, the angle between the vector from *s* to *s*^*t ′*^ (black bold line) and the the surface normal direction (red dashed arrow) is the cell tilt angle *ϕ* describing the degree of cell tilt at the local surface.

The cell density at the apical side is related to that at the basal side at *s* as *ρ*^*a*^(*s*) = *α*(*s*)*ρ*^*b*^(*s*). Since the total number of cells are the same at the two sides, the distribution of apico-to-basal ratio of density *α*(*s*) must follow:

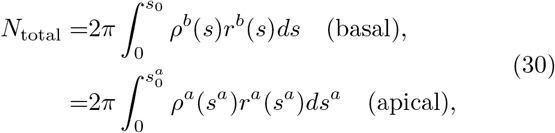

where the integration of *ds*^*a*^ (apical) or *ds* (basal) is over the whole meridional range *φ* ∈ [0, *π*] and 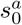 or *s*_0_ represents the half meridian and *N*_total_ is the total cell number covering the shell. When the apical element *dA*^*a*^ is only a normal projection of the basal element *dA* with a distance *e*(*s*), then one can obtain:

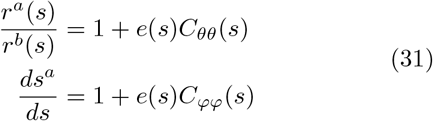

Inserting Eq. 31 into Eq. 29 leads to

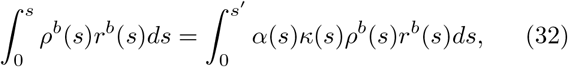

where *κ*(*s*) = 1 + 2*e*(*s*)*H*(*s*) + *e*^2^(*s*)*G*(*s*), with

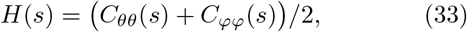

the mean curvature and

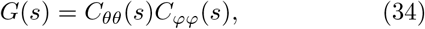

the Gaussian curvature.

Equation 32 now only depends on quantities with superscript *b*, so for neatness we omit this superscript from here and write *ρ*^*b*^(*s*) = *ρ*(*s*), *N*^*b*^(*s*) = *N* (*s*). Reorganizing the integration on the right hand side, we can transform Eq. 32 into

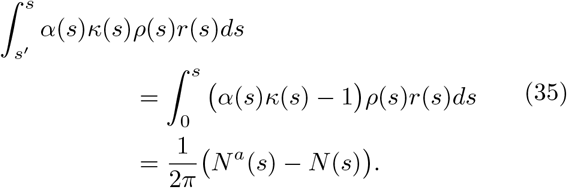

For *s*′ − *s* → 0, the left hand side of Eq. 35 is approximated as

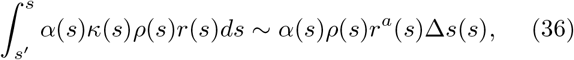

where Δ*s* is the arc length difference from *s* to *s*′ at the apical side. Accordingly, Eq. 35 becomes

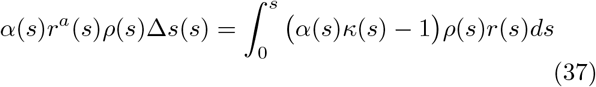

We then derive the tilt angle *ϕ* as:

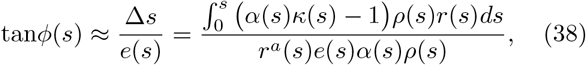

or in a more compact form

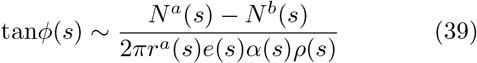

as shown in Eq. 4 in the main text.

Now we consider two extreme cases. If the the lateral membrane tension overwhelms the apical/basal layer tension, the lateral membranes tend to stand perpendicularly to the basal side. In this case, we have *α*(*s*) = 1*/κ*(*s*), so that *ϕ*(*s*) is zero across the space. By contrast, if the lateral membranes have low contractility compared with the apical/basal membranes, the cells tend to adjust the area sizes in both layers into homogeneous distributions, so that *α* becomes independent of the local curvature. We can derive a form for *α* as:

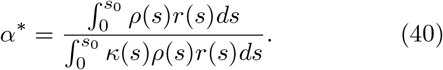

Inserting Eq. 40 into Eq. 38 gives

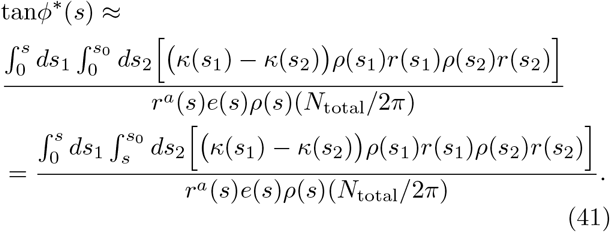

As the cell size is much smaller than the radius of curvature, *κ*(*s*_1_) − *κ*(*s*_2_) ∼ 2[*e*(*s*_1_)*H*(*s*_1_) − *e*(*s*_2_)*H*(*s*_2_)] with the second order term neglected. To clearly see the dependency of *ϕ*^*\**^ on curvature, we transform the integration of *ds* in Eq. 41 into integration by local cell number *dN* (*s*) = 2*πρ*(*s*)*r*(*s*)*ds* as

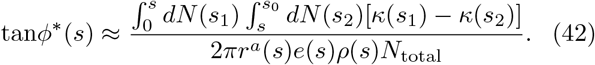

The integral 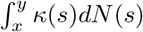 could be alternatively expressed as 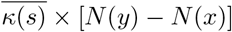, where

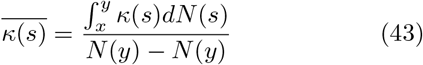

is the weighted average of *κ*(*s*) in the range of *x < s < y*. Hence, Eq. 42 be expressed as

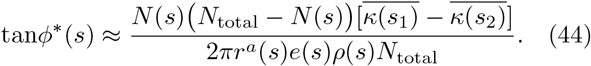

As long as the change of cell height *e*(*s*) with *s* is less radical than the change of curvature, we can approximate the difference of *κ* mainly by the change of mean curvature as 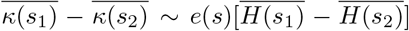, and finally we arrives at Eq. 6 in the main text.

We can relate the difference between the weighted average of the mean curvature to the mean curvature gradient. For 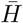 averaged from *x < s < y*, according to the integral mean value theorem, we can always find an 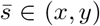 such that 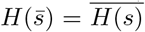. Hence, the difference of 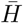could be re-expressed as

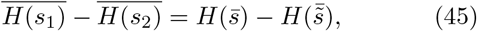

where 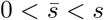 and 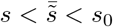. Since 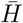 is the average weighted by the cell number at 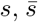 and 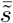 should be close to where the local cell number *dN* (*s*) is large, *i*.*e*., where *ρ*(*s*)*r*(*s*) is large. If the cell density does not radically change with *s, r*(*s*) will dominate where 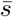 and 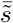 locates.

When *s* → 0 or)*s* → *s*_0_, *ϕ*^*\**^ is close to zero because *N* (*s*) (*N*_total_ − *N* (*s*)) → 0 and the contribution from the curvature becomes trivial. When *s* is neither close to the head (*s* = 0) nor the tail (*s* = *s*_0_), if the surface is convex, *r*(*s*) is large and 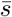 and 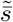 will be in a vicinity of *s*, where the distance 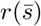 and 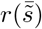 is relatively large. According to the mean value theorem, we could find another 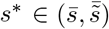 such that

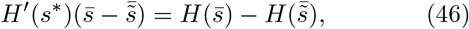

where *H*′ is the gradient of curvature and *s*^*\**^ is even more close to *s* than 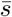 and 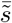. If the gradient of curvature is also continuous and differentiable (as seen in the ellipsoidal or tubular structures in biological systems), *H*′ (*s*^*\**^) ≈ *H*′ (*s*) + (*s*^*\**^*−s*)*H*^*″*^ (*s*); therefore, given a steep mean curvature gradient at *s*, we predict a large tilt angle *ϕ*^*\**^ at *s* in the zero-lateral-tension limit, as long as *s* is not at the head or the tail. Meanwhile, since 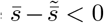, a negative gradient of curvature along *s* corresponds to the positive tilt angle towards the head. In other words, the tilt will lean to the vicinity of *s* with a higher positive curvature.

For more general cases, where the cells are subject to both the lateral and apical/basal layers, the distribution of *α*(*s*) is between 1*/κ*(*s*) and *α*^*\**^. Without other active sources, density projection rate *α*(*s*) together with the cell basal density *ρ*(*s*), and cell thickness *e*(*s*) are the mechanical consequence of cells minimizing their free energy as discussed further in Supp. Mat. G.

### E. Further simplifying the model for tilt

Using the mean value theorem to eliminate the integral Eq. 41, we obtain:

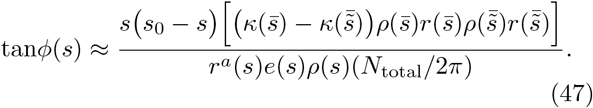

We can further simplify the model by setting a homogeneous density *ρ*(*s*) ∼*ρ*_0_ and *e*(*s*) ∼*ε* to arrive at a tilt profile purely depending on the geometry of the surface:

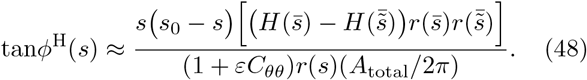

We can now evaluate the contributions from height modulation and basal density modulation separately. We define *ŝ* such that the total cell number at the apical side *N*_total_ = *ρ*(*ŝ*)*A*_total_, where *ρ*(*ŝ*) is a weighted average of density from *s* = 0 to *s* = *s*_0_. According to Eq. 48, the tilt profile with modulated inhomogeneous density becomes

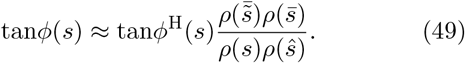

If *ρ*(*s*) is nearly homogeneous as |*dρ/ds*| ≪ 1, we assume *ρ*(*s*) ∼ *ρ*_0_[1 + *η*(*s*)(*s* − *ŝ*)*/s*_0_] with |*η*(*s*)| ≪ 1. Then,

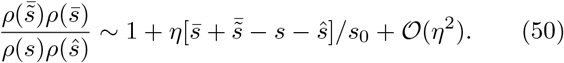

Recall that *ŝ* is the averaged position weighted by *ρ*(*s*)*r*(*s*) while 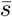 and 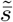 is the averaged position weighted by *κ*(*s*)*ρ*(*s*)*r*(*s*). With a radius of surface curvature much larger than the typical cell size - *κ* is only slightly larger than 1 (an assumption of our model) - we approximately have 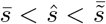. In this case, the first order term in *η* is negligible. In particular,

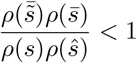

is valid when *ρ*(*s*) is a monotonic function. Therefore, the tilt angle under mild inhomogeneity of density is always slightly smaller the scenario with homogeneous density.

The tilt profile with modulated inhomogeneous cell height *e*(*s*) can be evaluated similarly, supposing *e*(*s*) = *ε*(1 + *η*′ (*s*)(*s*−*ŝ*)*/s*_0_) with |*η*′ (*s*) |≪1. Clearly, the value of *ε* has negligible effect on the result as long as *εH*(*s*) ≪1 (our basic model assumption) is valid.

Then, the tilt profile corrected by inhomogeneous cell height is

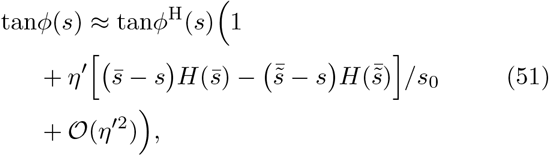

which has a more significant first order correction term in 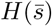 than Eq. 49. Therefore, inhomogeneity in cell height causes greater deviation of the tilt angle from the homogeneous limit *ϕ*^H^ than inhomogeneity in cell density.

In Fig. S4, we show the tilt profile and corresponding phase diagram for the AB-T1 transition measure for a prolate ellipsoidal system with *b/a* = 0.4 and *ε/a* = 0.05. The horizontal axis is the relative distance to the head and *d* = 1 represents the trunk. Tissue height and basal density are modulated linearly with *s*, with coefficients *β* and *λ* respectively. Modulation of density slightly suppresses the final tilt angle (the straight curves slightly lower than the dashed curves). Meanwhile, modulation of height affects the tilt more significantly, not only affecting the magnitude but also the shape of the distribution profile (as compared with the yellow lines, which corresponds to a homogeneous or zero modulation limit).

In conclusion, assuming the change of cell shape is relatively small to the change of curvature along the surface, we can simplify the model by ignoring the interdependency between the cell height and density. We just consider the inhomogeneity of cell height modulation, while keeping the density in either apical or basal side in a homogeneous setting.

**FIG. S4.**
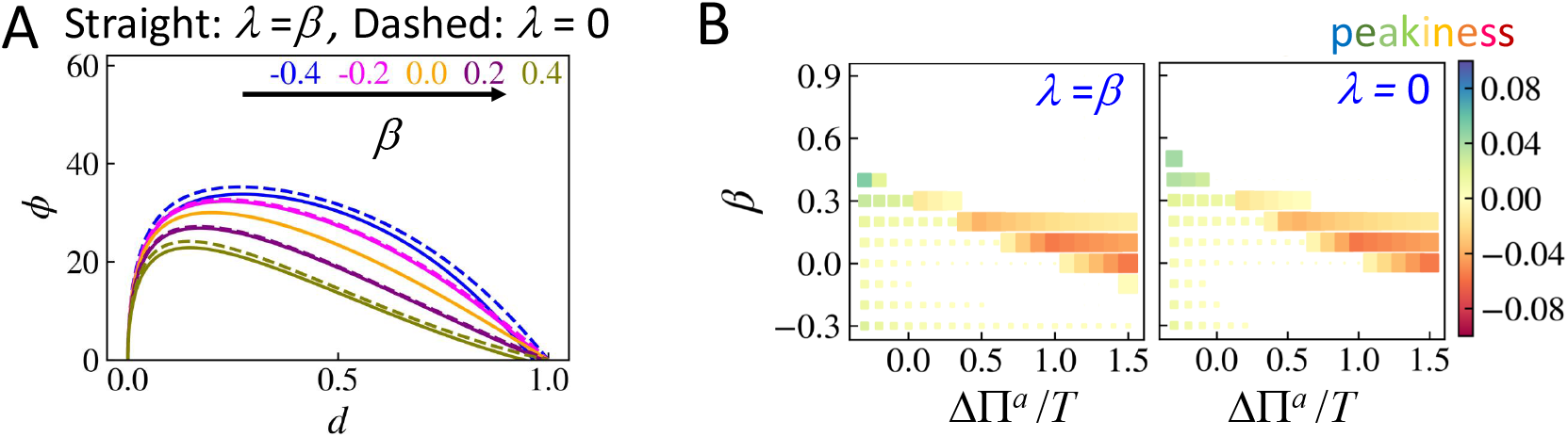
Results with an inhomogeneous basal density distribution modulated as *ρ*(*s*) = [*ρ*_0_ 1 + *λ*(*s/s*_1*/*4_ − 1*/*2)] and inhomogeneous cell height distribution *e*(*s*) = *ε* [1 + *β*(*s/s*_1*/*4_ 1*/*2)], where *s* is the arc length along the prime meridian surface and *s*_1*/*4_ is the 1*/*4 arc length. The system is a prolate with *b/a* = 0.4 and tissue height *ε/a* = 0.05. (A) A comparison of tilt angle profile between *λ* = *β* (straight) and *λ* = 0 (dashed) for varying *β* from −0.4 to 0.4. (B) A comparison of phase diagrams for the peak of the AB-T1 transition measure *γ* (main text Eq. 3). The color indicates the peak prominence, calculated as sign(*γ*_peak_) ×||*γ*_*peak*_| − |*γ*_trunk_||, and the size of the data square scales as ∝ (1−*d*_*peak*_)^2^ for a demonstration of the peak position. The closer the peak to the trunk, the smaller the data squares. If *d*_*peak*_ = 1 (peak at the trunk), the square is not visible.

### F. Tissue geometry on the surface of a prolate ellipsoid

Prolate spheroids were generated from revolving an ellipse around its the long axis as shown in Fig. S5A. An arbitrary point (*x, y, z*) on the surface of a prolate ellipsoid in 3D obeys

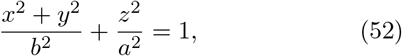

where *z* axis is the polar axis and 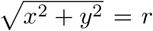 is the radial distance from the point to the polar axis *z*. Conventionally, *r* and *z* can be parameterized as

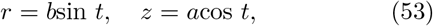

where *π/*2 − *t* is the reduced latitude of a spheroid. Considering elliptical symmetry, we discuss just the first quadrant (0 *< t < π/*2) in the following equations. The angle *φ* between the normal direction of a surface element and the polar axis is a function of *t* as

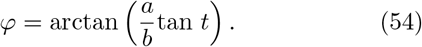

We assume the basal side of tissue is a surface of the prolate, while the apical side of the tissue is a projection on the normal direction with a distance

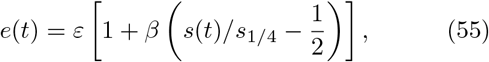

where *s*(*t*) is the arc length at *t, s*_1*/*4_ is 1/4 the ellipse perimeter and *β* is the rate of modulation of the tissue height. Note that *ds*(*t*) = *R*_*φφ*_(*t*)*dφ*(*t*).

The two principal curvatures of the surface element *dA*^*a*^(*t*) at the basal side are

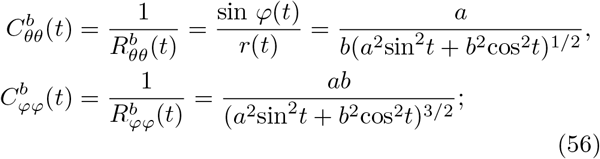

The curvature at the apical side depends on the tissue height *e*(*t*) as

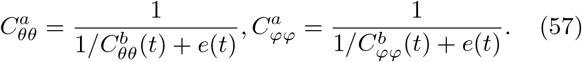

The projected area element at the apical side *dA*^*a*^(*t*) is larger than the area at the basal side *dA*^*b*^(*t*) by a ratio that decreases from the head of the prolate to the trunk due to the varying local principal curvatures. This area ratio can be expressed as:

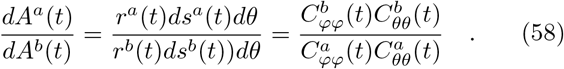

The horizontal axis *d* = 1 −cos(*t*) ∈ [0, 1] quantifies how close the point is to the head of the spheroid along the polar axis. For a smaller inverse aspect ratio *b/a*, the apico-to-basal area ratio is much larger at the head so that it decreases more sharply (straight curves for *b/a* = 0.3 and dashed curves for *b/a* = 0.4, Fig. S5). For *b* = *a*, the prolate ellipsoid is reduced to a sphere and the two principal curvatures become identical at any *t*, so that the apico-to-basal area ratio remains constant everywhere (inset in Fig. S5B). Meanwhile, a larger thickness of tissue causes a larger difference between the head and the trunk (different colors in Fig. S5B). The two stars in Fig. S5B indicate the area ratio measured from cells in the experiments (∼1.35 near the head and ∼1.23 near the trunk) for a relative thickness of tissue about 0.05 and an inverse aspect ratio *b/a* ∼ 0.4. Comparing these two points with the green dashed curve in Fig. S5B, we se that the cells at the apical surface, as measured in [29], are not perfectly normal projections of the basal layer. See Fig. S5C-D for the area ratios under inhomogeneous tissue height.

**FIG. S5.**
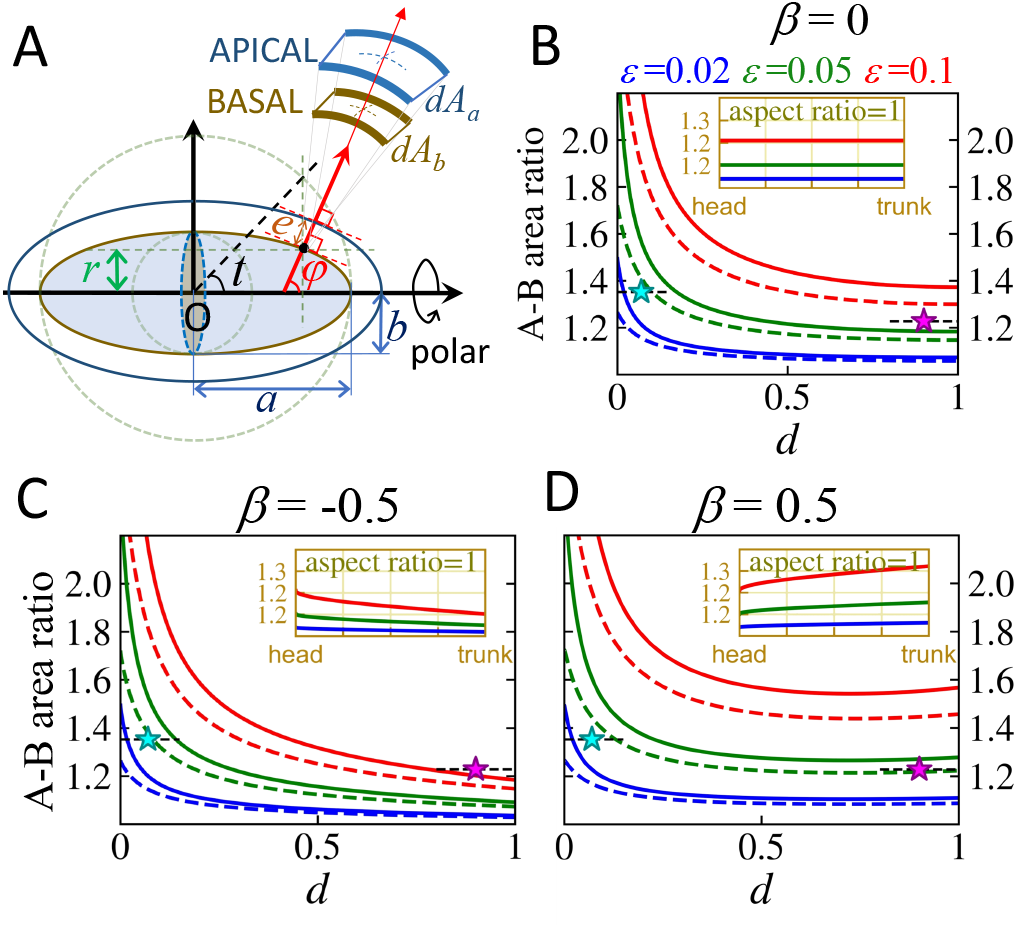
Apical to basal surface area on a prolate ellipsoid. (A) Illustration of the prolate ellipsoid. (B-D) Ratio of apical to basal area with different distance between the apical and_1_ the basal sides (*ϵ*) as a function of the arc length as *e*(*s*) =*ε*[1 + (*βs/s*_1/4_ − 1*/*2)], where *s*_0_ is the 1*/*4 of the perimeter of the meridian ellipse. (B) *β*=0; (C) *β <* 0; (D) *β >* 0. Different colors indicate different tissue heights *ε*. All the insets in (B-D) are for a spherical system (*a* = *b*). As a comparison, the two stars are the measured area ratios in the experimental data from [29] for wild type embryos; the ratio at the head is1 about 1.35 for the anterior side (*d <* 0.15) and the ratio at1 the trunk is about 1.23 for the trunk side (*d >* 0.7).

### G. Cell geometry control

The cell packing - which determines the cell areas on apical, basal and lateral sides - reaches a stable configuration when the system finds its minimal free energy. Here, we derive several analytical expressions for the tilt angle based on a mechanical model of cell geometry regulation. Following the established literature of vertex models [35, 43, 44], we describe the forces regulating th cell shape tissue as a derivative of the following free en ergy function:

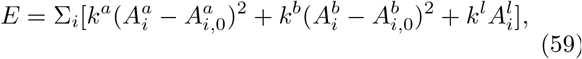

where 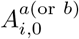 is the preferred cell area at the apical (or basal) layer for each cell *i. k*^*a*(or *b*)^ is the apical (or basal) elasticity coefficient and *k*^*l*^ is the lateral tension strength;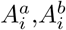 and 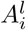are the areas of cell *i* at apical, basal and lateral surfaces respectively.

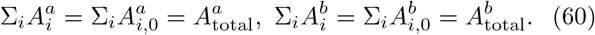

The packing equilibrium corresponds to the minimum of the free energy (Eq. 59) under the surface constraints given by Eq. 60.

If the lateral membrane is far less contractile than the apical and basal membranes, *i*.*e*., 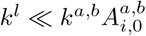, the minimization of this free energy will cause the cell to optimize its area towards 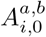in the apical and basal sides and the lateral membrane will tilt when the local curvatures of the shell change along the surface. By contrast, if 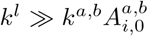, the minimization of the free energy leads to the lateral membrane orientating perpendicular to the apical and basal sides, with the cell apical area becoming a normal projection of the basal area, depending on the local curvatures. This can be seen from calculating the functional derivatives of Eq. 59.

For a demonstrative purpose, we show a derivation in a 2D equivalent and assume the preferred area of cells is homogeneous along the surface such that 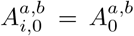 We first discuss a case without the single-cell volume constraint and then extend to a case with the volume constraint. The free energy in a 2D system is

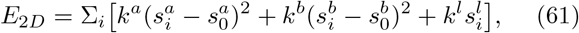

where 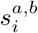 are the arc lengths of the cell at the apical or basal sides and 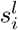 is the length of the cell lateral membrane. Note that cell height 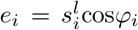, where *φ*_*i*_ is the tilt of lateral membrane of cell *i*. Minimizing this free energy constrains the cell side lengths so that the functional derivatives of the free energy become zero:

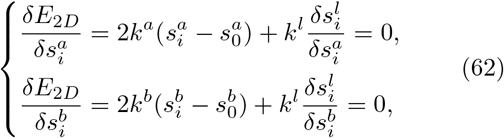

where 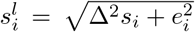as depicted by the line *ss*′ in Fig. S3. The surface constraint (Eq. 60) accordingly turns into a 1D form as

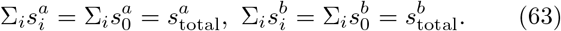

For simplicity, we assume *k*^*b*^→ ∞(due to the symmetry of the energy function, this assumption is equivalent to the case with a finite *k*^*b*^ but *k*^*a*^ → ∞), meaning that the basal layer is solid and thus the second equation in Eq. 62 can be ignored with 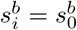.

Note that Δ*s*_*i*_ is also the difference between the 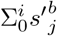, which is the accumulated normal projection length from the basal arc, and 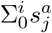, which is the accumulated apical arc length at cell *i*. Let *κ*_*i*_(*C*_*i*_, *e*_*i*_) be the normal projection rate merely depending on the curvature *C*_*i*_ and cell height *e*_*i*_ of cell *i*, and 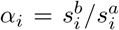be the ratio of basal length over apical length of cell *i* (in the large cell number limit, this quantity is equivalently defined before as the apico-to-basal ratio of cell density), then

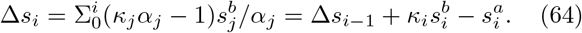

Eq. 62 then becomes

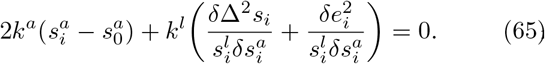

Without the volume constraint, the cell height distribution *e*_*i*_ is a consequence of multiple regulators, hence we take 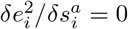. Eq. 65 reduces to:

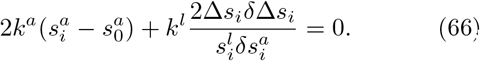

Inserting Eq. 64 into Eq. 66 we get

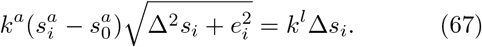

Let 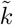be *k*^*l*^*/k*^*a*^*e*_*i*_. For an extreme limit 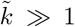 (rigid lateral membrane) we get

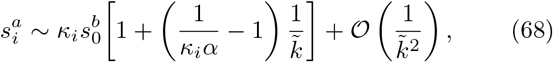

and for another extreme limit 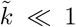 (less contractile lateral membrane) we get

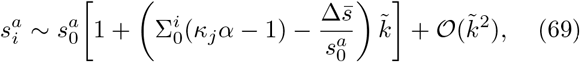

Where 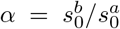, depending purely on the geometric information of the surfaces, and 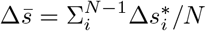 is the average tilted angle in a zero limit of lateral membrane contractility.

For a rigid membrane (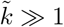, Eq. 68), the tilt angle is

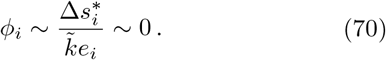

For a membrane with small contractility, (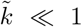, Eq. 69), substituting Eq. 69 into Eq. 64 leads to

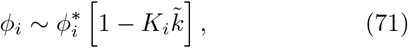

where 0 *< K*_*i*_∼1.

We next consider the effects of cell volume constraints. Now, the cell height, in relation to the basal and apical lengths, becomes

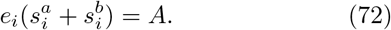

Inserting Eq. 72 into Eq. 65 and also assuming 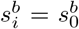, we obtain:

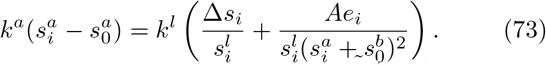

For a low contractility membrane, let 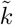 be *k*^*l*^*/k*^*a*^*ϵ*_*i*_ ≫ 1, then

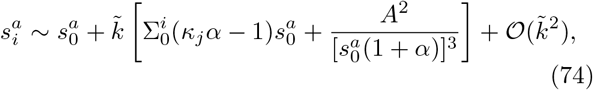

which has a correction term from the volume constraint *A* in the first order term of 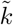as compared with Eq. 69. The tilt angle becomes

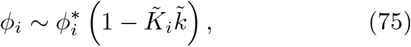

where 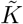 also has a correction ∼1 term from the volume constraint. Similarly, one can get the results under volume constraint for a rigid lateral membrane limit (not shown here).

